# Structure-based drug design and characterization of sulfonyl-piperazine benzothiazinone inhibitors of DprE1 from *Mycobacterium tuberculosis*

**DOI:** 10.1101/298539

**Authors:** Jérémie Piton, Anthony Vocat, Andréanne Lupien, Caroline Foo, Olga Riabova, Vadim Makarov, Stewart T. Cole

## Abstract

Macozinone (MCZ) is a tuberculosis (TB) drug candidate that specifically targets the essential flavoenzyme DprE1 thereby blocking synthesis of the cell wall precursor decaprenyl phosphoarabinose (DPA) and provoking lysis of *Mycobacterium tuberculosis*. As part of the MCZ back-up program we exploited structure-guided drug design to produce a new series of sulfone-containing derivatives, 2-sulphonylpiperazin 8-nitro 6-trifluoromethyl 1,3-benzothiazin-4-one, or sPBTZ. These compounds are less active than MCZ but have a better solubility profile and some derivatives display enhanced stability in microsomal assays. DprE1 was efficiently inhibited by sPBTZ and covalent adducts with the active site cysteine residue (C387) were formed. However, despite the H-bonding potential of the sulfone group no additional bonds were seen in the crystal structure of the sPBTZ-DprE1 complex with compound 11326127 as compared to MCZ. Compound 11626091, the most advanced sPBTZ, displayed good antitubercular activity in the murine model of chronic TB but was less effective than MCZ. Nonetheless, further testing of this MCZ backup compound is warranted as part of combination treatment with other TB drugs.

## INTRODUCTION

DprE1 is an essential flavoprotein of *Mycobacterium tuberculosis* involved in decaprenylphosphoryl-beta-D-arabinose (DPA) synthesis. DPA is the sole precursor of arabinose for production of both arabinogalactan and lipoarabinomannan (1), important components of the mycobacterial cell wall. DprE1, together with its counterpart DprE2, catalyse the epimerization of decaprenylphosphoryl-ß-D-ribose (DPR) to DPA in a two-step mechanism. In the last decade, many inhibitors were discovered to target DprE1, which is considered nowadays as the Achilles’ heel of *M. tuberculosis* due to its essentiality and especially to its localization in the periplasm (2). DprE1 inhibitors can be classified into two families according to their mode of action: some of them inhibit DprE1 irreversibly by forming a covalent adduct with cysteine 387 (C387) of DprE1, whereas others act as competitive non-covalent inhibitors (3).

The first covalent DprE1 inhibitors discovered were benzothiazinones (BTZ), exemplified by the lead compound, BTZ043, which is exceptionally potent with *in vitro* and *ex vivo* minimal inhibitory concentration (MIC) values in the nanomolar range (4). A lead optimization campaign gave rise to PBTZ169, now known as Macozinone (MCZ). It is currently the most potent BTZ compound against *M. tuberculosis* with an MIC of 0.3 nM (5), has completed preclinical development successfully and is now undergoing Phase I and phase II clinical trials (https://www.newtbdrugs.org/pipeline/clinical). A common characteristic of the covalent DprE1 inhibitors is the presence of a nitro group on the molecule, which is essential for the mechanism of inhibition. Indeed, this nitro group is converted by DprE1 containing FADH_2_ into an extremely reactive nitroso group which specifically targets the cysteine residue at position 387 (C387) in the active site of DprE1, to form a covalent adduct and thereby irreversibly inhibits the enzyme (4–7, 8, 9–13)}. It has been demonstrated that the presence of C387 is essential for the activity of covalent DprE1 inhibitors (4, 14).

Apart from the covalent bond with C387, covalent DprE1 inhibitors are otherwise only maintained in the pocket by steric hindrance and Van Der Walls interactions, which explains why a simple substitution at C387 results in complete resistance of the enzyme to these compounds (3). As part of the MCZ-back-up program, this observation prompted us to revisit the Structure Activity Relationship (SAR) in order to obtain derivatives with other anchor points in the active site of DprE1, since such compounds might retain activity against C387 mutants should these arise.

Several non-covalent DprE1 inhibitors have also been found (15-24). Similar to the covalent DprE1 inhibitors, these non-covalent compounds sit in the substrate-binding pocket of DprE1 and act as competitive inhibitors. Interestingly, a class of non-covalent inhibitors, 2-carboxyquinoxalines, are active against BTZ-resistant *M. tuberculosis* strains with substitutions at C387 of DprE1 (14). Molecules from this family possess an essential 2-carboxylate moiety that forms key hydrogen bonds with the side-chain of Lysine 418 and the hydroxyl group of Tyrosine 60 (18). Hence, it can be hypothesised that a composite molecule between 2-carboxyquinoxalines and MCZ could overcome resistance issues and increase specificity to the target.

Revisiting the SAR of MCZ further provides the opportunity to improve pharmacodynamics properties of MCZ such as aqueous solubility to increase its oral bioavailability, metabolic stability, and *in vivo* activity (25). It has been observed that the potency of BTZ derivatives is inversely proportional to their solubility (4, 5). Therefore, the solubility and bioavailability of the most active benzothiazinones are parameters for improvement. Factors controlling the aqueous solubility of organic molecules are complex and drug solubility issues are usually solved by a combination of empirical and rational drug design strategies. More than twenty crystal structures of DprE1 with or without inhibitors were reviewed recently to identify the structural determinants for activity and to guide rational drug design (3).

Based on our prior observations, the aim of this study was to design a new structure-guided series of MCZ derivatives with increased activity against either wild-type *M. tuberculosis* or its BTZ-resistant mutants, and with improved solubility, absorption, bioavailability and metabolic stability of the compound *in vivo*. Therefore, a new series of MCZ derivatives, harbouring a sulfonylpiperazine group, was designed (2-sulphonylpiperazin 8-nitro 6-trifluoromethyl 1,3-benzothiazin-4-one or sPBTZ), synthesized and characterized. This study identifies 11626091 as the best sPBTZ and demonstrates that this compound has a promising combination of antitubercular activity and ADME/T properties.

## RESULTS

### Rationale

When BTZ inhibitors bind to their target, DprE1, the sole bond formed is a covalent semimercaptal bond with the active site cysteine residue, C387. We reasoned first that introducing a sulfonyl group into the MCZ scaffold might offer a second anchor by mimicking the carboxylate moiety of the 2-carboxyquinoxalines that acts as a H-bond acceptor with DprE1 and increases affinity of the inhibitor for the target. Secondly, sulfonyl groups are well characterized and present in many FDA-approved drugs, in particular anti-mycobacterial agents, such as dapsone. Based on these observations, sulfonyl groups might increase both the solubility and bioavailability of the inhibitor *in vivo*. Lastly, the geometry imposed by sulfonyl groups opens new directions for investigation of the SAR.

### Synthesis of sPBTZ

The synthesis of 17 sulfanyl-piperazino BTZ (sPBTZ) derivatives was performed in a two-step procedure from 2-(methylthio)-8-nitro-6-(trifluoromethyl)-4*H*-1,3-benzothiazin-4-one, as described previously (Makarov et al., 2014). Its reaction with a 5-molar excess of free piperazine generated the corresponding piperazine derivative with a high yield. This scaffold was used in the reactions with different alkyl-, aryl-or heteryl-sulfochlorides to form sulfanyl-piperazino BTZs. The compounds synthesized have different types of sulfonyl substitutions thus allowing the structure-activity relationship to be studied. It is clear that compounds with alkyl substitutions have much better antitubercular activity and aryl derivatives have much lower activity, consistent with our previous data for piperazine-containing BTZ (PBTZ) derivatives (Makarov et al., 2014).

### Solubility

The octanol-water partition coefficient, logP, which is regarded as a suitable indicator of molecular hydrophobicity and bioavailability, was calculated for all derivatives to measure the effects of introducing a sulfonyl group on the PBTZ backbone. Interestingly, the addition of the sulfonyl group between the benzothiazinone and piperazine moieties has the tendency to decrease the clogP coefficient and therefore hydrophobicity (Table 1). The sulfonylated derivative of MCZ, sPBTZ169 (11326127) that carries a sulfonyl group between the piperazine and cyclohexyl moieties, has a clogP of 3.28 whereas MCZ has a clogP of 4.31. This implies that the introduction of a sulfonyl group decreases hydrophobicity, and may thus increase solubility in physiological conditions and subsequently could have an important impact on bioavailability.

**Table 1:**
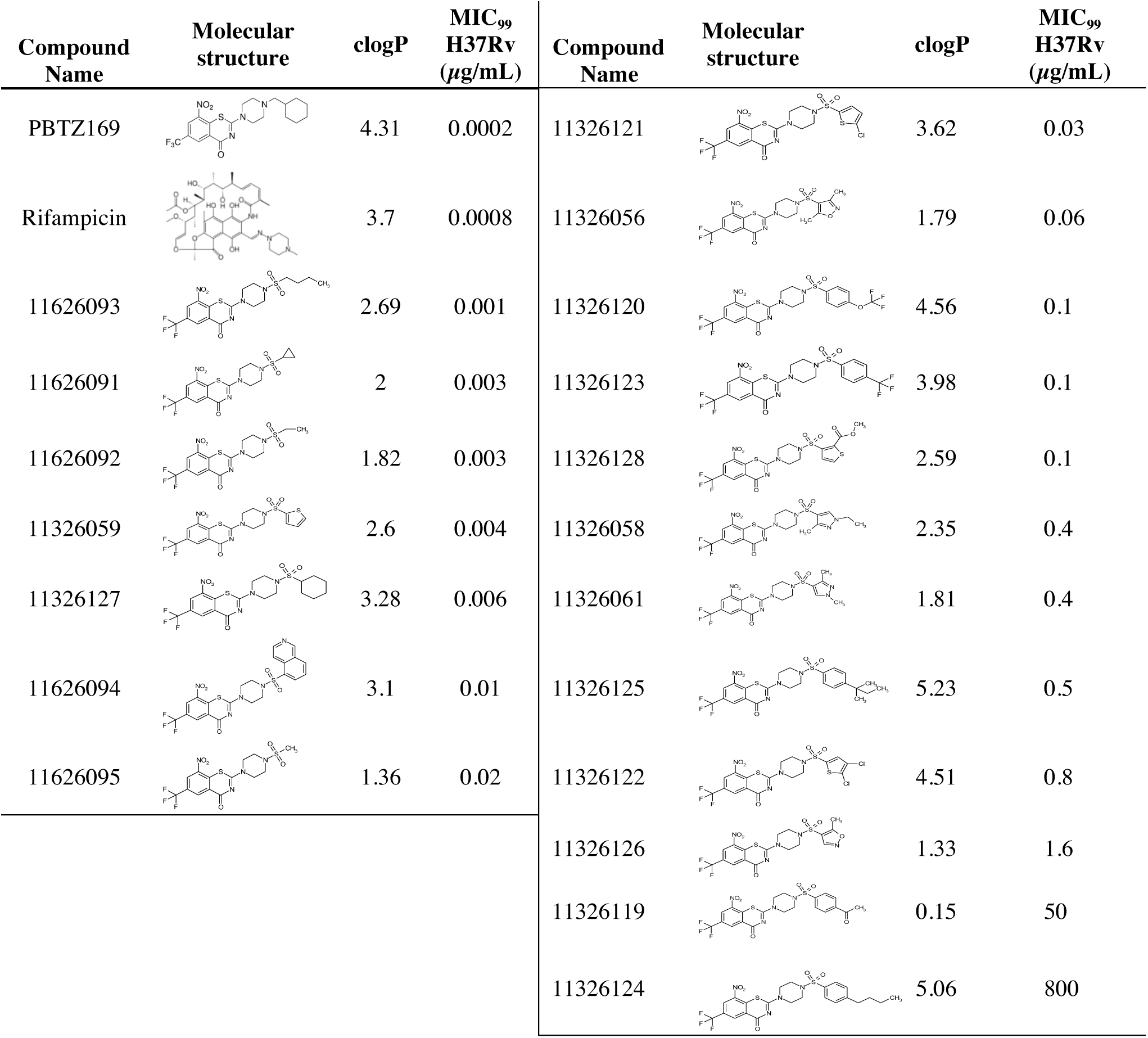
Structure activity relationship of the different 2-sulphonylpiperazin 8-nitro 6-trifluoromethyl 1,3-benzothiazin-4-ones derivatives (sulfonyl PBTZ derivatives) in *M. tuberculosis* H37Rv

| Compound Name | Molecular structure | clogP | MIC<sub>99</sub> H37Rv (µg/mL) | Compound Name | Molecular structure | clogP | MIC<sub>99</sub> H37Rv (µg/mL) |
| --- | --- | --- | --- | --- | --- | --- | --- |
| PBTZ169 |  | 4.31 | 0.0002 | 11326121 |  | 3.62 | 0.03 |
| Rifampicin |  | 3.7 | 0.0008 | 11326056 |  | 1.79 | 0.06 |
| 11626093 |  | 2.69 | 0.001 | 11326120 |  | 4.56 | 0.1 |
| 11626091 |  | 2 | 0.003 | 11326123 |  | 3.98 | 0.1 |
| 11626092 |  | 1.82 | 0.003 | 11326128 |  | 2.59 | 0.1 |
| 11326059 |  | 2.6 | 0.004 | 11326058 |  | 2.35 | 0.4 |
| 11326127 |  | 3.28 | 0.006 | 11326061 |  | 1.81 | 0.4 |
| 11626094 |  | 3.1 | 0.01 | 11326125 |  | 5.23 | 0.5 |
| 11626095 |  | 1.36 | 0.02 | 11326122 |  | 4.51 | 0.8 |
|  |  |  |  | 11326126 |  | 1.33 | 1.6 |
|  |  |  |  | 11326119 |  | 0.15 | 50 |
|  |  |  |  | 11326124 |  | 5.06 | 800 |

### Antitubercular activity

The activity of all sPBTZ derivatives was tested *in vitro* against *M. tuberculosis* H37Rv and MIC_99_ values were determined (Table 1). Most sPBTZ were active *in vitro* in the sub-micromolar range, proving that that addition of the sulfonyl group does not abolish activity. However, none has a better activity than MCZ. 11626093, which has a butyl group, has the highest activity of MIC 1 ng/*μ*L corresponding to 5 times the MIC_99_ of MCZ. The introduction of a sulfone between the piperazine and the cyclohexyl negatively affected the activity of the compound, reducing activity by 30 fold, as observed with sPBTZ169 (11326127), which has an MIC_99_ of 6 nM. It was previously shown that substituting the methylcyclohexyl in MCZ with small radicals such as methyl or ethyl decreases the activity of compounds, with MIC of 250 ng/mL and 60 ng/mL, respectively (5). Interestingly, when methylcyclohexyl is substituted by sulfonylmethyl (11626095) or sulfonylethyl (11626092), the activity decreases less and the compounds are 10-times more active that the non-sulfonated derivatives (MIC of 20 ng/mL and 3 ng/mL, respectively). Substitution with a butyl leads to the same activity independent of the sulfonyl group. This observation indicates that the presence of the sulfonyl group positively affects activity when the radical is small (methyl or ethyl) whereas it negatively influences activity when the substituent is long and hydrophobic.

Similarly to the other BTZ derivatives, the activities of the sulfonyl derivatives are inversely proportional to the logP (4), suggesting that activity could be related to solubility in lipids (Figure S1). Seven molecules were selected based on their activity/logP profile for further characterisation (Figure S1).

### Target engagement and structural studies

To ensure that sPBTZs specifically target DprE1, selected sulfonyl derivatives were tested against *M. tuberculosis* NTB1, a DprE1 mutant that carries a cysteine 387 serine (C387S) substitution and is thus resistant to BTZ. As expected, none of the sBTZs are active against NTB1, indicating that DprE1 is their only target (Table 2). Furthermore, it is evident that the sulfonyl group is unable to mimick the carboxylate moiety of 2-carboxyquinoxalines in stabilising the compound in the pocket as a non-covalent inhibitor.

**Table 2:** *In vitro* activity of selected sPBTZ derivatives against *M. tuberculosis* H37Rv and its BTZ-resistant mutant (NTB1) and cytotoxicity for HepG2 cells

| Compound<br>Name | H37Rv<br>MIC<sub>99</sub><br>(µg/mL) | NTB1<br>MIC<sub>99</sub><br>(µg/mL) | TD50<br>(µg/mL) | Selective<br>index |
| --- | --- | --- | --- | --- |
| 11326059 | 0.004 | >100 | 100 | 25000 |
| 11326127 | 0.006 | >100 | >100 | >16666 |
| 11626091 | 0.003 | 14.7 | 8.10 | 2700 |
| 11626092 | 0.003 | 25.1 | 8.90 | 2967 |
| 11626093 | 0.001 | >100 | >100 | >100000 |
| 11626094 | 0.01 | >100 | 100.00 | 10000 |
| 11626095 | 0.02 | 75 | 7.80 | 390 |
| PBTZ169 | 0.0002 | >100 | 44.00 | 220000 |
| Rifampicin | 0.0008 | 0.0008 | 100.00 | 125000 |

To investigate whether the introduction of the sulfonyl group could influence the position of the inhibitor and allow more contact within the binding pocket, a crystal structure of DprE1 in complex with sPBTZ169 was solved and compared to the crystal structure of DprE1 with MCZ. sPBTZ169 is located in the same pocket as MCZ and other BTZ derivatives. It sits in an hydrophobic pocket *via* the trifluoromethyl group and is covalently bound to C387 (Figure 1). The compound is maintained by Van der Walls interaction on each side by V365, Y314, W230 and FAD, and a hydrogen bond between K418 and the oxygen atom of the nitro group of sPBTZ169. Unfortunately, the orientation of sPBTZ does not favor the formation of a new anchor to the protein for instance interaction between the sulfonyl function and Y60. Futhermore as MCZ, the electron density map does not account fully for the sulfonyl-cyclohexyl moiety of sPBTZ169 likely due to its higher flexibility demonstrating that sulfonyl is not stabilized in the pocket ((5)).

**Figure 1:**
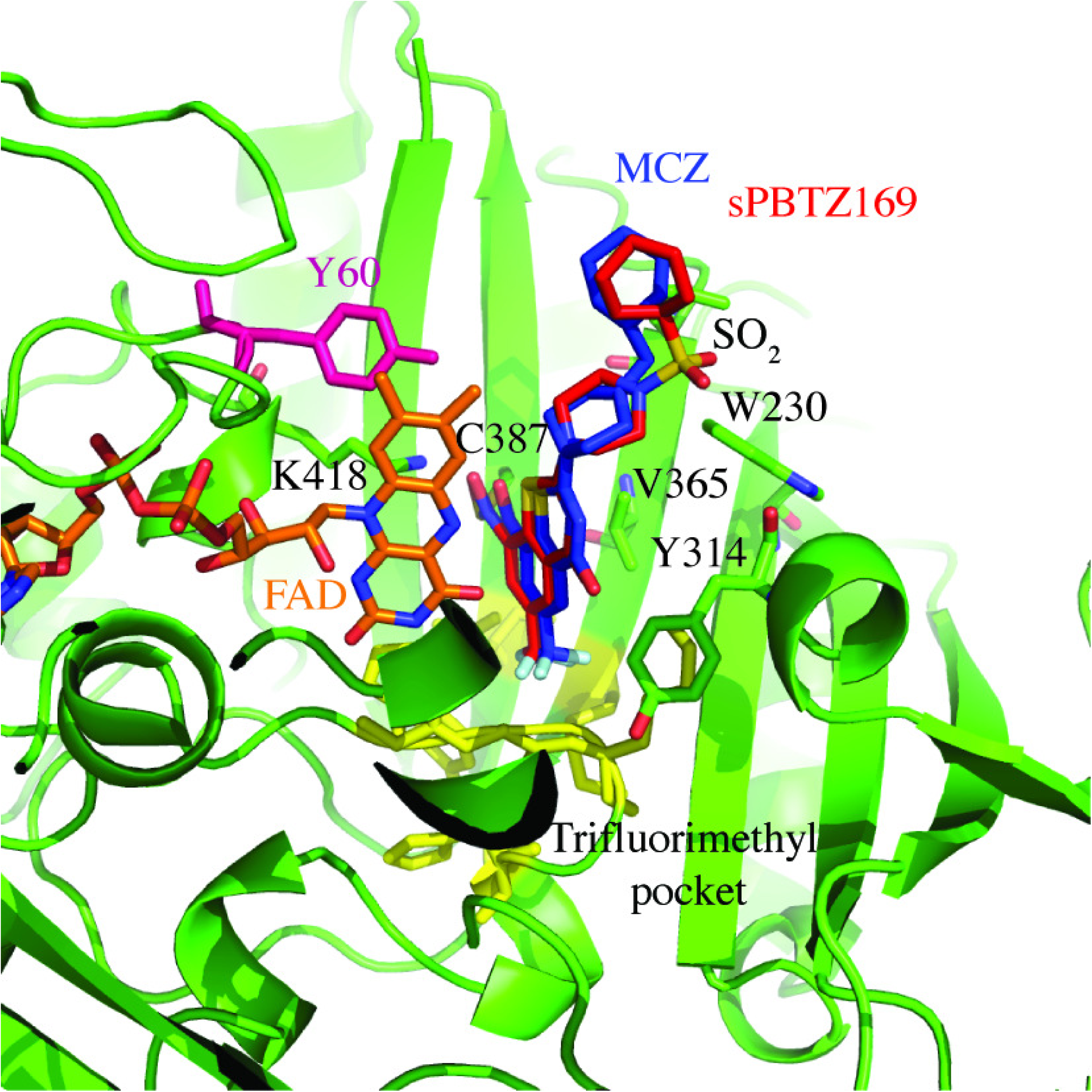
Structural comparison between the crystal structure of DprE1 in complex with sulfonyl derivative sPBTZ169 (11326127) and the structure in complex with MCZ. (PDB code: 4NCR (5)). DprE1 is represented in green in the cartoon. sPBTZ169 (in red) sits in the trifluoromethyl hydrophobic pocket (in yellow) and binds covalently to C387. It is maintained by FAD (orange) and some lateral chains represented in sticks. Y60, a key residue in the binding of 2-Carboxyquinoxalines, is represented in pink.

To determine whether the sulfonyl group could affect the activity at the protein level and even help to stabilize the inhibitor in the active site of a BTZ-resistant C387S DprE1 variant, the inhibition of DprE1 activity was measured *in vitro*. IC_50_ values were determined for the wild type and the BTZ-resistant C387S mutant enzyme for MCZ and sPBTZ169 (Table S2). sPBTZ169 has an IC_50_ of 1.1 and 12 *μ* M against wild type and BTZ-resistant DprE1, respectively, whereas the corresponding IC_50_ values for MCZ are 0.3 and 3.6 *μ*M. That introducing the sulfonyl group on the MCZ scaffold leads to 4-fold less activity suggests that even if the environment of the protein is favorable for an H-bond acceptor, the presence of a hydrophobic group, such as a cyclohexyl in MCZ, is still preferable for activity of the drug. On the other hand, the higher IC_50_ against C387S compared to the WT enzyme validates the structural studies in that the sulfonyl group does not help to stabilize the molecule in the pocket of the BTZ-resistant mutant C387S.

### ADME/T

To assess potential cytotoxic effects of the sulfonyl group on the sPBTZ derivatives, viability of HepG2 cells was monitored after exposure to different concentrations of the compounds. The concentration for half-maximal cytotoxicity (TD_50_) was determined for each compound. Four of the seven compounds were not cytotoxic (11326059, 11326127, 11626093, 1162694) while three of them showed mild cytotoxicity at concentrations of around 10 *μ*g/mL (1162691, 1162692, 1162695). Taken together, the selective index representing the ratio of the antitubercular activity of compounds (MIC_99_ on H37Rv) to cytotoxicity (TD_50_ on HepG2 cells) is more than acceptable for the chosen seven compounds.

Since solubility issues are often encountered in drug development, which would consequently impact bioavailability and activity *in vivo*, the solubility of these derivatives was calculated by the shake flask method in equilibrium in water. Experimental solubility constants measured were then compared to theoretical solubility constants calculated using the SwissADME webserver (26). sPBTZ are predicted to have increased solubility compared to MCZ in water (Table 3) and experimental solubility constants measured in water for all sPBTZ derivatives are consistent with theoretical constants calculated by SwissADME using different algorithms (Table 3). As expected, 11626095, 11626092 and 11626091, harboring methyl, ethyl and cyclopropyl groups, respectively, are the most soluble sPBTZ derivatives. In contrast, 11326059, 11326127, 11626093, and 11626094 that carry bigger hydrophobic groups are at least 100-times less soluble in water. Interestingly, there is a good correlation between water solubility and clogP. However, a discrepancy between the predicted and experimental solubility for MCZ was observed. In fact, MCZ is 400 times more soluble in water than calculated with different algorithms.

**Table 3:** Water solubility of selected sPBTZ derivatives measured by shake flask method compared to solubility calculated on SwissAdme ((26))

| **Compound**<br>**Name** | **Solubility measured**<br>**in water ($\mu\text{g/mL}$)** | **Solubility calculated**<br>**in water ($\mu\text{g/ml}$)** | **clogP** |
| --- | --- | --- | --- |
| 11326059 | <0.01 | 0.12 | 2.6 |
| 11326127 | <0.01 | 0.26 | 3.28 |
| 11626091 | $8.6 \pm 3.8$ | 4.69 | 1.91 |
| 11626092 | $10.7 \pm 4.6$ | 7.2 | 2 |
| 11626093 | $0.105 \pm 0.05$ | 0.93 | 2.69 |
| 11626094 | $0.088 \pm 0.04$ | 0.18 | 3.1 |
| 11626095 | $10.6 \pm 1.8$ | 16.9 | 1.36 |
| PBTZ169 | $31.1 \pm 6.4$ | 0.0795 | 4.31 |

**Table 4:** Metabolic stability of sPBTZ derivatives measured in mouse and human microsomes

| Compound Name | Cl<sub>int</sub> mouse<br>microsome<br>(µl/min/mg protein) | Cl<sub>int</sub> human microsome<br>(µl/min/mg protein) | References |
| --- | --- | --- | --- |
| 11326059 | 18.6 | 16.8 | This study |
| 11326127 | 19 | 21.5 | This study |
| 11626091 | 2.7 | 9.7 | This study |
| 11626092 | 3 | 1.4 | This study |
| 11626093 | 42 | 6.4 | This study |
| 11626094 | 24 | 277 | This study |
| 11626095 | 0.9 | 1.0 | This study |
| Carbamazepine | 0.6 | 0.4 | This study |
| Nifedipine | 105 | 121.1 | This study |
| PBTZ169 | 36.7 | 28 | (5) |
| BTZ043 | 16.5 | 4.20 | (5) |

Another possible issue in the development of MCZ might be metabolic instability (5). In order to test if the sulfonyl group could improve metabolic stability effected by mouse or human liver enzymes, microsomal stability experiments were conducted. Compounds 11626095, 11626091 and 11626092 harbouring the smallest radical chains methyl, ethyl and cyclopropyl, respectively, have the lowest clearances in both mouse and human microsomes, indicative of their stability. 11626093 carrying a butyl radical is metabolically stable in human but highly unstable in mouse microsomes, suggesting that it could be a good substrate for mouse but not human microsomal enzymes. Compounds 11326059 and 11326127 have medium clearances, similar to MCZ and BTZ043. Of note, 11626094 is highly unstable in the presence of human microsomes. These results reveal a strong correlation between the length of the substituent after the piperazine moiety and instability in microsomes.

### Activity in the murine model of chronic TB

Finally, the *in vivo* efficacy of 1162691 was assessed in the murine model of chronic TB following lowdose aerosol infection of BALB/cByJ mice with *M. tuberculosis* and treatment at 50 mg/kg (Figure 2). Compound 1162691 was selected as it is the most promising taking into account its activity (“only” 15 times less active than MCZ), cytotoxicity, solubility, and metabolic stability properties.

**Figure 2:**
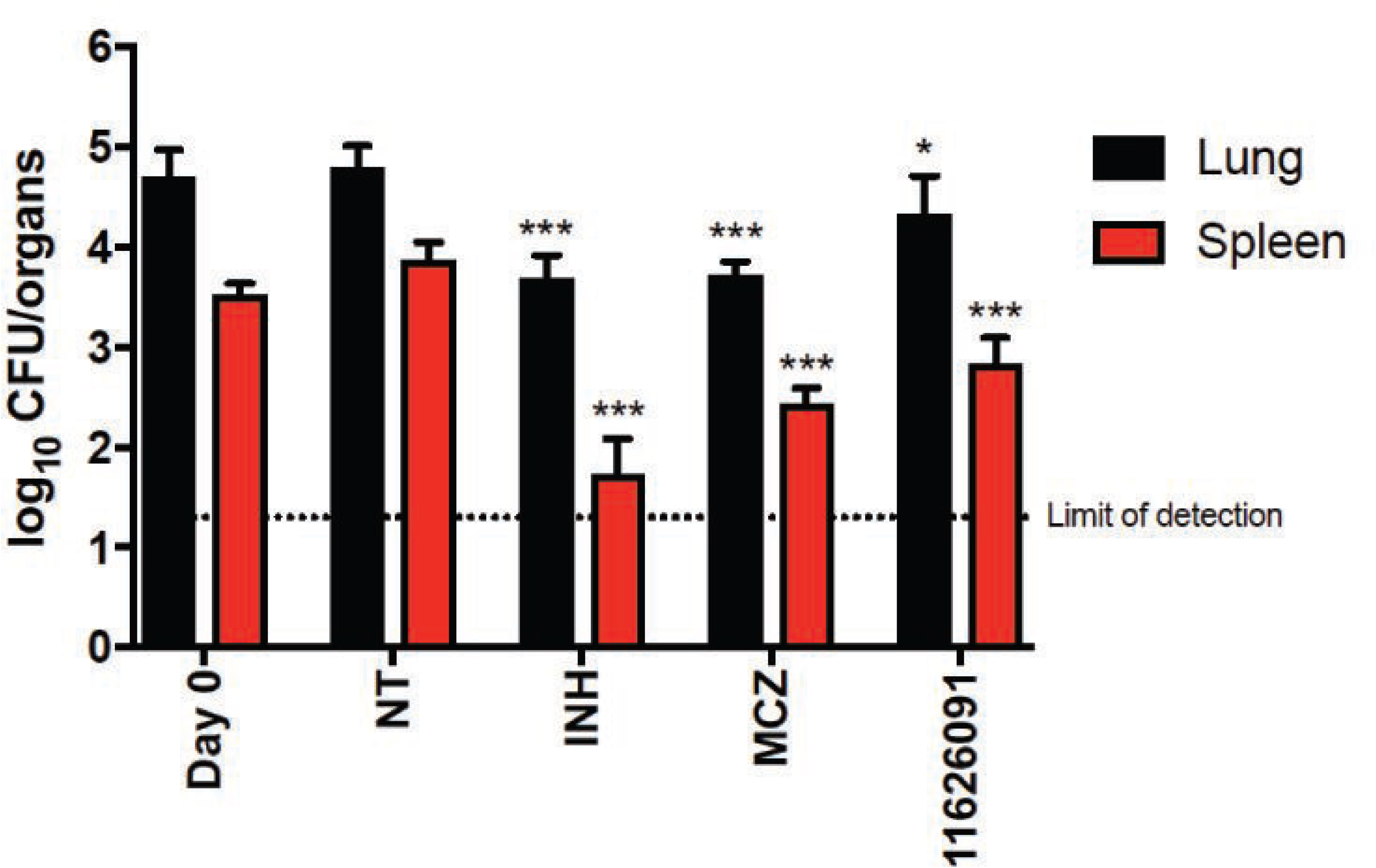
Activity of the sPBTZ derivative 11626091 (50 mg/kg) compared to *in vivo* activity of INH (10 mg/kg) or MCZ (25 mg/kg) in the mouse model of chronic TB. Black columns correspond to the bacterial burden in lungs and red columns correspond to the bacterial burden in the spleens, at day 0 (D0) when treatment was initiated, or day 28 (NT) when treatment ended. Bars represent the mean ± s.d. of CFUs from 5 mice per group. NT: untreated control. Limit of detection: 20 CFU/organ. The significance of difference was calculated using the Student t-test. *P<0.05. **P<0.005. ***P<0.0001 compared to NT.

Compared to the untreated control group, the bacterial burden in the lungs and spleens of 1162691-treated mice was 0.46 (*) and 1.03 (***) log_10_ CFU lower, respectively (Fig 2), whereas the values for those organs from MCZ-treated mice were 1.03 (***) and 1.47 (***) log_10_ CFU. These results indicate that 1162691 is highly active *in vivo* in the murine model of chronic TB, although it is less able to reduce the bacterial load in the lungs than treatment with MCZ. Interestingly, 1162691 seems to be more efficacious in the spleen than in lungs.

## DISCUSSION

Lead compounds for new anti-TB drugs should possess not only potent sterilizing activity but also good pharmacokinetic profiles to facilitate their co-administration with other anti-TB and anti-HIV (human immunodeficiency virus) agents. New drugs should also be appropriate for once daily oral dosing and be relatively inexpensive to produce to ensure that all high-burden countries have access. Structure-based rational drug design supports drug development and has the potential to increase the activity and pharmacokinetic properties of lead compounds.

The new sPBTZ series was designed as part of the MCZ back-up program. MCZ is a BCS (biopharmaceutical classification system) class 2 drug with a low dissolution rate but excellent absorption (5). Structural studies indicate that the environment of the binding pocket in the protein can accommodate a polar group at the cyclohexyl position of MCZ (3). A sulfonyl group was deemed to be a good candidate because it is well characterized and present in many soluble and metabolically stable FDA-approved drugs including those for treating mycobacterial infections. For instance, sulfamethoxazole is a sulfonamide drug used in co-trimoxazole prophylaxis for HIV-infected patients, demonstrating that it is compatible with anti-retroviral treatment (27).

It was previously shown that there is a strong correlation between logP and BTZ activity (4). We found that introduction of a sulfonyl group into the MCZ scaffold to form the sPBTZ series decreases both logP and activity *in vitro* (Table 1). However, sPBTZ still retain potency and the presence of the sulfonyl group improved aqueous solubility for those derivatives, which harbour small side chains, such as methyl, ethyl or cyclopropyl, as compared to MCZ (5). Interestingly, aqueous solubility and metabolic stability measured in microsomes indicate a better behaviour for sPBTZ with small groups rather than those with long hydrophobic chains or MCZ. Despite being less active than MCZ our sulfonylated PBTZ with long hydrophobic chains, methyl (11626095), ethyl (11626092) or cyclopropyl (11626091) derivatives are considered as good candidates in terms of their solubility and metabolic stability profiles. The efficacy of the most active derivative of the three, 11626091, was assessed *in vivo* in the murine model of chronic TB, where relatively good activity was measured in the lungs and particularly in the spleens (Figure 2). It is also important to note that dissolution of 11626091 was considerably easier than MCZ in methylcellulose, the solvent used for the *in vivo* studies (technical observation, data not shown).

One of the objectives of inserting a sulfonyl group into the PBTZ backbone was to increase mitigate potential BTZ-resistance by increasing the number of hydrogen bonds with DprE1. Indeed, a sulfonyl could act as an H-bond acceptor in order to anchor the protein to the H-bond donors, for example the hydroxyl group of Tyrosine 60 localized in the binding pocket. This group was identified as a key player in the stabilization of 2-carboxyquinoxalines, molecules that remain active against BTZ-resistant DprE1. Unfortunately, as was observed in the crystal structure and enzymatic inhibition assays, the sulfonyl group of sPBTZ is not implicated in the binding and stabilization of the drug in the pocket as originally hypothesized. This explains the resistance of the BTZ-resistant *M. tuberculosis* mutant NTB1 to sPBTZ.

To conclude, our study identifies 11626091 as an active, metabolically stable and moderately soluble molecule that is less active than MCZ *in vitro* and *in vivo*. However, given its better solubility, compound 11626091 represents an attractive back-up to MCZ, that should now be tested in combination with other TB drug candidates such as bedaquiline in order to assess its potential to contribute to a new regimen.

## MATERIALS AND METHODS

### Synthesis

The synthetic route used to produce sPBTZ and related procedures are described in the supporting materials.

### Octanol-water partition coefficient logP calculation

The log*P* values were calculated using the program Hyperchem 7.5 (Hypercube Inc., http://www.hyper.com).

### Bacterial strains and culture conditions

*M. tuberculosis* strain H37Rv and its BTZ-resistant mutant (C387S) NTB1 were grown at 37°C with shaking in Middlebrook 7H9 broth (Difco) supplemented with 10% albumin-dextrose-catalase (ADC) enrichment, 0.2% glycerol, and 0.05% Tween 80. The *in vitro* activities against all mycobacterial strains were measured with the resazurin reduction microplate assay (REMA), by 2-fold serial dilutions of the compounds in the working bacterial culture in 96-well plates (final volume of 100 *μ*l). Following incubation for 6 days at 37°C bacterial viability was determined by adding resazurin for 24 h at 37°C and measuring the fluorescence of the resorufin metabolite (excitation wavelength, 560 nm; emission wavelength, 590 nm) using a Tecan Infinite M200 microplate reader. Briefly, the noise signal obtained in the blank control wells was subtracted from the fluorescence values of test samples and mycobacterial viability in each well were proportionally calculated compared to 100% growth in control wells. Bacterial viability curves and MIC_99_ value were calculated with GraphPad Prism software version 7.0, using “Gompertz equation for MIC determination” analysis (GraphPad Software, Inc., La Jolla, CA, USA).

### Cytotoxicity studies

The cytotoxicity of the compounds was measured as described previously against the human hepatic cell line, HepG2, (20). Briefly, cells were incubated (4,000 cells/well) with serial dilutions of compounds (2-fold dilutions; 100 to 0.1 *μ*g/ml) in a 96-well microplate. Following incubation for 3 days at 37°C cell viability was determined by adding resazurin for 4 h at 37°C and measuring the fluorescence of the resorufin metabolite (excitation wavelength, 560 nm; emission wavelength, 590 nm) using a Tecan Infinite M200 microplate reader. Data were corrected for background (no-cell control) and expressed as a percentage of the value for untreated cells (cells only).

Data were fitted to obtain IC_50_ values, using “Log(inhibitor) vs. response - Variable slope” function implemented in GraphPad Prism software version 7.0 (GraphPad Software, Inc., La Jolla, CA, USA). Selective index refers to the ratio of the dose of drug that causes toxicity effects (TD50) to the dose that leads to the desired pharmacological effect (MIC_99_).

### Water Solubility

Water solubility measurements were performed using the shake flask method ((28)). Approximately 1 mL was added to 1 mg of compound in an Eppendorf tube and incubated for 3 days at 25°C with shaking at 800 rpm. Suspensions were centrifuged at 16,100 x g for 10 min and the supernatant filtered using 0.22 *μ* filters. Filtrates were injected onto a high-performance liquid chromatography (HPLC) column (Dionex) and the amount of compound quantified using a calibration curve.

### Inhibition assays, crystallography and structural studies of DprE1 complexed with sPBTZ. Recombinant

*M. tuberculosis* DprE1 was overexpressed and purified as described previously (5) to obtain highly concentrated and pure protein with bound flavin adenine dinucleotide (FAD). Enzyme activities in the presence of MCZ, Ty38c and 11326127 were measured as described previously to determine IC_50_ (50% inhibitory concentrations) values for both wild-type and C131S mutant DprE1 proteins (14).

For crystallization purposes, *M. tuberculosis* DprE1 (approximately 40 *μ* M) was incubated for 3h at 30 °C, with 200 *μ* M sulfonyl-BTZ 11326127 and 200 *μ* M FPR (farnesyl phosphoribose), in 20 mM Tris pH 8.5, 50 mM NaCl. The protein was concentrated to approximately 15 mg/mL on an Amicon centrifugal device (30,000 MWCO, Millipore). Crystals were obtained by the hanging-drop vapor diffusion method at 18 °C. Experiments were set up by mixing 1 *μ* 1 of the protein sample with 1 *μ*1 of the reservoir solution containing 100 mM imidazole, pH 7.2-7.5, 18-24 % polypropyleneglycol 400. Yellow crystals grew in approximately 1-3 days and were transferred to a cryo-protectant (reservoir solution with 25% glycerol) prior to flash-cooling in liquid nitrogen.

X-ray data were collected at SLS, beamline PROXIMA 3. Data were integrated with the program XDS (29) and processed using PHENIX (30). Structure was solved by molecular replacement using PHENIX (31) and the structure of *M. tuberculosis* DprE1 (PDB code : 4NCR, (5)) as a template. Molecular replacement was subjected to iterative rounds of refinement and rebuilding in coot (32) and PHENIX. The coordinates and structure factors have been deposited in the Brookhaven Protein Data Bank (accession numbers 6G83).

### Metabolic stability in vitro

The intrinsic clearance (CL_int_) of compounds was measured in both mouse and human liver microsomes. Briefly, 100 *μ*g of mouse (CD-1) or human liver microsomes (both from Invitrogen) were mixed in 0.1M phosphate buffer (pH 7.4) containing 0.01 *μ*l of compound dissolved in DMSO at 10 mg/ml, in a final volume of 50 *μ*l. In parallel, a NADPH-regenerating system (Promega) was prepared in 0.1 M phosphate buffer (pH 7.4). The solutions were pre-incubated at 37°C for 10 min before the intrinsic clearance assessment was initiated by mixing the two solutions (50 *μ*l of each; final compound concentration, 1 *μ*g/ml) at 37°C. After 0, 5, 15, 30, and 60 min, the reactions were terminated by transferring 100 *μ*l of the reaction mixture into 100 *μ*l of acetonitrile and placing the mixture on ice for 30 min for full protein precipitation. Samples were then centrifuged at 12,000 x g for 10 min, and the supernatant was injected onto a high-performance liquid chromatography (HPLC) column (Dionex) to quantify the amount of parent compound remaining over time. Carbamazepine and Nifedipine at the same concentrations were used as controls for low and high intrinsic clearances respectively.

### Anti-mycobacterial activity of 11626091 against chronic TB in mice

Female BALB/cByJ mice, aged 5 to 6 weeks, were purchased from Charles River Laboratories (France). Mice were infected with a low dose aerosol (100 - 200 CFU/lung) of logarithmic-phase *M. tuberculosis* H37Rv bacilli, then were allocated to experimental groups and returned to their cages. Five mice were used per time point for each regimen. Treatment was initiated 4 weeks after infection.

Macozinone (MCZ), 11626091, and isoniazid (INH) were prepared weekly in 0.5% methylcellulose and administered at 25, 50, and 10 mg/kg, respectively, by gavage 5 days a week for 4 weeks. *In vivo* efficacy of each treatment was assessed by CFU enumeration after plating dilutions of the lung and spleen homogenates on 7H10 agar plates containing 10% OADC, cycloheximide (10 μg/ml), and ampicillin (50 μg/ml). Plates were incubated for 4 weeks at 37°C before CFU were enumerated. CFU counts were log10 transformed before analysis as mean log10 CFU ± standard deviation (SD), and were compared by Student’s t-test using GraphPad Prism^®^ version 7.0 software (GraphPad Software, Inc., La Jolla, CA, USA). P values less than 0.05 were considered as statistically significant.

Experiments were approved by the Swiss Cantonal Veterinary Authority (authorization no. 3082) and performed between June-August 2017.

## ACKNOWLEDGMENTS

The research leading to these results was performed as part of the More Medicines for Tuberculosis (MM4TB) project and received funding from the European Community’s Seventh Framework Programme ([FP7/ 2007-2013]) under grant agreement no. 260872. We thank Aline Reynaud and Florence Pojer from the Ppscf facility at EPFL for their technical support.

## REFERENCES

1. Mikusova K, Huang H, Yagi T, Holsters M, Vereecke D, D’Haeze W, Scherman MS, Brennan PJ, McNeil MR, Crick DC. 2005. Decaprenylphosphoryl arabinofuranose, the donor of the D-arabinofuranosyl residues of mycobacterial arabinan, is formed via a two-step epimerization of decaprenylphosphoryl ribose. Journal of bacteriology 187:8020–5.

2. Brecik M, Centarova I, Mukherjee R, Kolly GS, Huszar S, Bobovska A, Kilacskova E, Mokosova V, Svetlikova Z, Sarkan M, Neres J, Kordulakova J, Cole ST, Mikusova K. 2015. DprE1 Is a Vulnerable Tuberculosis Drug Target Due to Its Cell Wall Localization. ACS Chem Biol 10:1631–6.

3. Piton J, Foo CS, Cole ST. 2016. Structural studies of Mycobacterium tuberculosis DprE1 interacting with its inhibitors. Drug Discov Today doi:10.1016/j.drudis.2016.09.014.

4. Makarov V, Manina G, Mikusova K, Mollmann U, Ryabova O, Saint-Joanis B, Dhar N, Pasca MR, Buroni S, Lucarelli AP, Milano A, De Rossi E, Belanova M, Bobovska A, Dianiskova P, Kordulakova J, Sala C, Fullam E, Schneider P, McKinney JD, Brodin P, Christophe T, Waddell S, Butcher P, Albrethsen J, Rosenkrands I, Brosch R, Nandi V, Bharath S, Gaonkar S, Shandil RK, Balasubramanian V, Balganesh T, Tyagi S, Grosset J, Riccardi G, Cole ST. 2009. Benzothiazinones kill Mycobacterium tuberculosis by blocking arabinan synthesis. Science 324:801–4.

5. Makarov V, Lechartier B, Zhang M, Neres J, van der Sar AM, Raadsen SA, Hartkoorn RC, Ryabova OB, Vocat A, Decosterd LA, Widmer N, Buclin T, Bitter W, Andries K, Pojer F, Dyson PJ, Cole ST. 2014. Towards a new combination therapy for tuberculosis with next generation benzothiazinones. EMBO Mol Med 6:372–83.

6. Batt SM, Jabeen T, Bhowruth V, Quill L, Lund PA, Eggeling L, Alderwick LJ, Futterer K, Besra GS. 2012. Structural basis of inhibition of Mycobacterium tuberculosis DprE1 by benzothiazinone inhibitors. Proc Natl Acad Sci U S A 109:11354–9.

7. Trefzer C, Rengifo-Gonzalez M, Hinner MJ, Schneider P, Makarov V, Cole ST, Johnsson K. 2010. Benzothiazinones: prodrugs that covalently modify the decaprenylphosphoryl-beta-D-ribose 2’-epimerase DprE1 of Mycobacterium tuberculosis. J Am Chem Soc 132:13663–5.

8. Neres J, Pojer F, Molteni E, Chiarelli LR, Dhar N, Boy-Rottger S, Buroni S, Fullam E, Degiacomi G, Lucarelli AP, Read RJ, Zanoni G, Edmondson DE, De Rossi E, Pasca MR, McKinney JD, Dyson PJ, Riccardi G, Mattevi A, Cole ST, Binda C. 2012. Structural basis for benzothiazinone-mediated killing of Mycobacterium tuberculosis. Science translational medicine 4:150ra121.

9. Christophe T, Jackson M, Jeon HK, Fenistein D, Contreras-Dominguez M, Kim J, Genovesio A, Carralot JP, Ewann F, Kim EH, Lee SY, Kang S, Seo MJ, Park EJ, Skovierova H, Pham H, Riccardi G, Nam JY, Marsollier L, Kempf M, Joly-Guillou ML, Oh T, Shin WK, No Z, Nehrbass U, Brosch R, Cole ST, Brodin P. 2009. High content screening identifies decaprenyl-phosphoribose 2’ epimerase as a target for intracellular antimycobacterial inhibitors. PLoS Pathog 5:e1000645.

10. Magnet S, Hartkoorn RC, Szekely R, Pato J, Triccas JA, Schneider P, Szantai-Kis C, Orfi L, Chambon M, Banfi D, Bueno M, Turcatti G, Keri G, Cole ST. 2010. Leads for antitubercular compounds from kinase inhibitor library screens. Tuberculosis (Edinb) 90:354–60.

11. Stanley SA, Grant SS, Kawate T, Iwase N, Shimizu M, Wivagg C, Silvis M, Kazyanskaya E, Aquadro J, Golas A, Fitzgerald M, Dai H, Zhang L, Hung DT. 2012. Identification of novel inhibitors of M. tuberculosis growth using whole cell based high-throughput screening. ACS Chem Biol 7:1377–84.

12. Landge S, Mullick AB, Nagalapur K, Neres J, Subbulakshmi V, Murugan K, Ghosh A, Sadler C, Fellows MD, Humnabadkar V, Mahadevaswamy J, Vachaspati P, Sharma S, Kaur P, Mallya M, Rudrapatna S, Awasthy D, Sambandamurthy VK, Pojer F, Cole ST, Balganesh TS, Ugarkar BG, Balasubramanian V, Bandodkar BS, Panda M, Ramachandran V. 2015. Discovery of benzothiazoles as antimycobacterial agents: Synthesis, structure-activity relationships and binding studies with Mycobacterium tuberculosis decaprenylphosphoryl-beta-D-ribose 2’-oxidase. Bioorg Med Chem 23:7694–710.

13. Landge S, Ramachandran V, Kumar A, Neres J, Murugan K, Sadler C, Fellows MD, Humnabadkar V, Vachaspati P, Raichurkar A, Sharma S, Ravishankar S, Guptha S, Sambandamurthy VK, Balganesh TS, Ugarkar BG, Balasubramanian V, Bandodkar BS, Panda M. 2016. Nitroarenes as Antitubercular Agents: Stereoelectronic Modulation to Mitigate Mutagenicity. ChemMedChem 11:331–9.

14. Foo CS, Lechartier B, Kolly GS, Boy-Rottger S, Neres J, Rybniker J, Lupien A, Sala C, Piton J, Cole ST. 2016. Characterization of DprE1-Mediated Benzothiazinone Resistance in Mycobacterium tuberculosis. Antimicrob Agents Chemother 60:6451–6459.

15. Wang F, Sambandan D, Halder R, Wang J, Batt SM, Weinrick B, Ahmad I, Yang P, Zhang Y, Kim J, Hassani M, Huszar S, Trefzer C, Ma Z, Kaneko T, Mdluli KE, Franzblau S, Chatterjee AK, Johnsson K, Mikusova K, Besra GS, Futterer K, Robbins SH, Barnes SW, Walker JR, Jacobs WR, Jr., Schultz PG. 2013. Identification of a small molecule with activity against drug-resistant and persistent tuberculosis. Proc Natl Acad Sci U S A 110:E2510–7.

16. Shirude PS, Shandil R, Sadler C, Naik M, Hosagrahara V, Hameed S, Shinde V, Bathula C, Humnabadkar V, Kumar N, Reddy J, Panduga V, Sharma S, Ambady A, Hegde N, Whiteaker J, McLaughlin RE, Gardner H, Madhavapeddi P, Ramachandran V, Kaur P, Narayan A, Guptha S, Awasthy D, Narayan C, Mahadevaswamy J, Vishwas KG, Ahuja V, Srivastava A, Prabhakar KR, Bharath S, Kale R, Ramaiah M, Choudhury NR, Sambandamurthy VK, Solapure S, Iyer PS, Narayanan S, Chatterji M. 2013. Azaindoles: noncovalent DprE1 inhibitors from scaffold morphing efforts, kill Mycobacterium tuberculosis and are efficacious in vivo. J Med Chem 56:9701–8.

17. Chatterji M, Shandil R, Manjunatha MR, Solapure S, Ramachandran V, Kumar N, Saralaya R, Panduga V, Reddy J, Prabhakar KR, Sharma S, Sadler C, Cooper CB, Mdluli K, Iyer PS, Narayanan S, Shirude PS. 2014. 1,4-azaindole, a potential drug candidate for treatment of tuberculosis. Antimicrob Agents Chemother 58:5325–31.

18. Neres J, Hartkoorn RC, Chiarelli LR, Gadupudi R, Pasca MR, Mori G, Venturelli A, Savina S, Makarov V, Kolly GS, Molteni E, Binda C, Dhar N, Ferrari S, Brodin P, Delorme V, Landry V, de Jesus Lopes Ribeiro AL, Farina D, Saxena P, Pojer F, Carta A, Luciani R, Porta A, Zanoni G, De Rossi E, Costi MP, Riccardi G, Cole ST. 2015. 2-Carboxyquinoxalines kill mycobacterium tuberculosis through noncovalent inhibition of DprE1. ACS Chem Biol 10:705–14.

19. Naik M, Humnabadkar V, Tantry SJ, Panda M, Narayan A, Guptha S, Panduga V, Manjrekar P, Jena LK, Koushik K, Shanbhag G, Jatheendranath S, Manjunatha MR, Gorai G, Bathula C, Rudrapatna S, Achar V, Sharma S, Ambady A, Hegde N, Mahadevaswamy J, Kaur P, Sambandamurthy VK, Awasthy D, Narayan C, Ravishankar S, Madhavapeddi P, Reddy J, Prabhakar K, Saralaya R, Chatterji M, Whiteaker J, McLaughlin B, Chiarelli LR, Riccardi G, Pasca MR, Binda C, Neres J, Dhar N, Signorino-Gelo F, McKinney JD, Ramachandran V, Shandil R, Tommasi R, Iyer PS, Narayanan S, Hosagrahara V, Kavanagh S, Dinesh N, Ghorpade SR. 2014. 4-aminoquinolone piperidine amides: noncovalent inhibitors of DprE1 with long residence time and potent antimycobacterial activity. J Med Chem 57:5419–34.

20. Makarov V, Neres J, Hartkoorn RC, Ryabova OB, Kazakova E, Sarkan M, Huszar S, Piton J, Kolly GS, Vocat A, Conroy TM, Mikusova K, Cole ST. 2015. The 8-Pyrrole-Benzothiazinones Are Noncovalent Inhibitors of DprE1 from Mycobacterium tuberculosis. Antimicrob Agents Chemother 59:4446–52.

21. Shaikh MH, Subhedar DD, Arkile M, Khedkar VM, Jadhav N, Sarkar D, Shingate BB. 2016. Synthesis and bioactivity of novel triazole incorporated benzothiazinone derivatives as antitubercular and antioxidant agent. Bioorg Med Chem Lett 26:561–9.

22. Tiwari R, Miller PA, Chiarelli LR, Mori G, Sarkan M, Centarova I, Cho S, Mikusova K, Franzblau SG, Oliver AG, Miller MJ. 2016. Design, Syntheses, and Anti-TB Activity of 1,3-Benzothiazinone Azide and Click Chemistry Products Inspired by BTZ043. ACS Med Chem Lett 7:266–70.

23. Panda M, Ramachandran S, Ramachandran V, Shirude PS, Humnabadkar V, Nagalapur K, Sharma S, Kaur P, Guptha S, Narayan A, Mahadevaswamy J, Ambady A, Hegde N, Rudrapatna SS, Hosagrahara VP, Sambandamurthy VK, Raichurkar A. 2014. Discovery of pyrazolopyridones as a novel class of noncovalent DprE1 inhibitor with potent anti-mycobacterial activity. J Med Chem 57:4761–71.

24. Chikhale R, Menghani S, Babu R, Bansode R, Bhargavi G, Karodia N, Rajasekharan MV, Paradkar A, Khedekar P. 2015. Development of selective DprE1 inhibitors: Design, synthesis, crystal structure and antitubercular activity of benzothiazolylpyrimidine-5-carboxamides. Eur J Med Chem 96:30–46.

25. Amidon GL, Lennernas H, Shah VP, Crison JR. 1995. A theoretical basis for a biopharmaceutic drug classification: the correlation of in vitro drug product dissolution and in vivo bioavailability. Pharm Res 12:413–20.

26. Daina A, Michielin O, Zoete V. 2017. SwissADME: a free web tool to evaluate pharmacokinetics, drug-likeness and medicinal chemistry friendliness of small molecules. Sci Rep 7:42717.

27. Guidelines W. 2014. Guidelines on Post-Exposure Prophylaxis for HIV and the Use of Co-Trimoxazole Prophylaxis for HIV-Related Infections Among Adults, Adolescents and Children: Recommendations for a Public Health Approach: December 2014 supplement to the 2013 consolidated guidelines on the use of antiretroviral drugs for treating and preventing HIV infection, Geneva.

28. Bergstrom CA, Wassvik CM, Johansson K, Hubatsch I. 2007. Poorly soluble marketed drugs display solvation limited solubility. J Med Chem 50:5858–62.

29. Kabsch W. 2010. Integration, scaling, space-group assignment and post-refinement. Acta Crystallogr D Biol Crystallogr 66:133–44.

30. Adams PD, Afonine PV, Bunkoczi G, Chen VB, Davis IW, Echols N, Headd JJ, Hung LW, Kapral GJ, Grosse-Kunstleve RW, McCoy AJ, Moriarty NW, Oeffner R, Read RJ, Richardson DC, Richardson JS, Terwilliger TC, Zwart PH. 2010. PHENIX: a comprehensive Python-based system for macromolecular structure solution. Acta Crystallogr D Biol Crystallogr 66:213–21.

31. McCoy AJ, Grosse-Kunstleve RW, Adams PD, Winn MD, Storoni LC, Read RJ. 2007. Phaser crystallographic software. J Appl Crystallogr 40:658–674.

32. Emsley P, Cowtan K. 2004. Coot: model-building tools for molecular graphics. Acta Crystallogr D Biol Crystallogr 60:2126–32.

